# Molecular and mineral responses of corals grown under artificial Calcite Sea conditions

**DOI:** 10.1101/2022.02.25.481970

**Authors:** Nicola Conci, Erika Griesshaber, Ramón E. Rivera-Vicéns, Wolfgang W. Schmahl, Sergio Vargas, Gert Wörheide

## Abstract

The formation of skeletal structures composed of different calcium carbonate polymorphs (aragonite and calcite) appears to be regulated both biologically and environmentally. Among environmental factors influencing aragonite and calcite precipitation, changes in seawater conditions – primarily in the molar ratio of magnesium and calcium during so-called “Calcite” (*m*Mg:*m*Ca below 2) or “Aragonite” seas (*m*Mg:*m*Ca above 2) – have had profound impacts on the distribution and performance of marine calcifiers throughout the Earth’s history. Nonetheless, the fossil record shows that some species appear to have counteracted such changes and kept their skeleton polymorph unaltered. Here, the aragonitic octocoral *Heliopora coerulea* and the aragonitic scleractinian *Montipora digitata* were exposed to Calcite Sea-like *m*Mg:*m*Ca with various levels of changes in magnesium and calcium concentration, and both mineralogical (i.e., CaCO_3_ polymorph) and gene expression changes were monitored. Both species maintained aragonite deposition at lower *m*Mg:*m*Ca ratios, while concurrent calcite presence was only detected in *M. digitata*. Despite a strong variability between independent experimental replicates for both species, the expression for a set of putative calcification-related genes, including known components of scleractinian skeleton organic matrix, was found to consistently change at lower *m*Mg:*m*Ca. These results support previously proposed involvements of the skeleton organic matrix in counteracting decreases in seawater *m*Mg:*m*Ca. Although no consistent changes in expression for calcium and magnesium transporters were observed, down-regulation calcium channels in *H. coerulea* in one experimental replicate and at an *m*Mg:*m*Ca of 2.5 might indicate the possibility of active calcium uptake regulation by the corals under altered *m*Mg:*m*Ca.

## Introduction

Biomineralization refers to the process by which organisms secrete and form their skeletons (Lowenstam and Weiner, 1989). Anthozoan (Phylum Cnidaria) biomineralization holds a particularly important ecological role, as it represents the primary mechanism sustaining the growth of coral reefs. In corals, skeletons are composed of different calcium carbonate (CaCO_3_) polymorphs. Members of the order Scleractinia produce aragonite exoskeletons, except *Paraconotrochus antarcticus*, a recently discovered species that exhibits a two-component (i.e., aragonite and calcite) skeleton (Stolarski et al., 2021). In the subclass Octocorallia, an opposite pattern occurs: the vast majority of species produce calcitic endoskeletal structures, while aragonite exoskeletons are deposited by members of the order Helioporacea (i.e., the blue corals).

As observed in other organisms, including mollusks, arthropods, and chordates, biomineralization in corals is a *biologically controlled* process (Lowenstam and Weiner, 1989). During biomineralization, corals control the availability and concentration of ions required for calcification (e.g., Ca^2+^ and HCO_3_ ^-^) in the extracellular calcifying medium (ECM) (Moya et al., 2008; Bertucci et al., 2011; Zoccola et al., 2015; Le Goff et al., 2016), and the production of various macromolecules putatively involved in calcium carbonate deposition, collectively referred to as the skeleton organic matrix (SkOM). SkOM components are secreted into the calcification space and are eventually occluded within the skeleton mineral. Several regulatory functions have been attributed to the SkOM of corals and other marine calcifiers. These include crystal nucleation, formation, and inhibition (Wheeler et al., 1981; Peled-Kamar et al., 2002; Puverel et al., 2005; Von Euw et al., 2017), and controlling the calcium carbonate (CaCO_3_) polymorph of the skeleton (Thompson et al., 2000; Goffredo et al., 2011).

In corals, the effect of different organic compounds on CaCO_3_ polymorphs (aragonite, calcite) has been investigated mostly through *in vitro* precipitation experiments. The addition of total SkOM extracts (Hohn and Reymond, 2019), or proteins (Rahman and Oomori, 2009; Goffredo et al., 2011) and lipids (Reggi et al., 2016) isolated from coral skeletons to supersaturated CaCO_3_ solutions, has been shown to promote or inhibit the formation of specific CaCO_3_ polymorphs and to affect the shape of CaCO_3_ crystals. Recently, Laipnik et al. (2019) observed that protein-driven *in vitro* precipitation of different CaCO_3_ polymorphs is also related to the magnesium concentration ([Mg^2+^]) in the solution used. In those experiments, the absence of aragonite formation – the naturally occurring polymorph in scleractinian skeletons – at low magnesium concentrations led the authors to propose a key role for seawater [Mg^2+^] in the functioning of coral skeletogenic proteins.

Seawater chemistry has in fact been long included among the main drivers of selective inorganic precipitation of different CaCO_3_ polymorphs, with the molar ratio of magnesium and calcium (*m*Mg:*m*Ca) likely representing one of the key factors leading to differential CaCO_3_ precipitation (Morse and Mackenzie, 1990; Morse et al., 1997; Balthasar and Cusack, 2015). From an evolutionary perspective, the *m*Mg:*m*Ca is particularly interesting due to its estimated fluctuations in the last 600 million years, which caused alternating periods of calcite- and aragonite-favoring oceanic environments (Sandberg, 1983) that robustly correlate with the preferred CaCO_3_ skeleton polymorph of the dominant reef builders during those geological periods (Stanley and Hardie, 1998, 1999). During the Cretaceous, for example, when the *m*Mg:*m*Ca dropped to ∼ 1 (compared to the modern value of 5.2), scleractinian corals were replaced by calcitic bivalves (rudists, Phylum Mollusca) as main reef builders. Remarkably, some calcitic scleractinians have been reported from this period <u>(Stolarski et al., 2007)</u>. However, fossils of cretaceous aragonitic corals indicate that some coral species could deposit aragonite despite the lower, calcite-favoring *m*Mg:*m*Ca marine environment (Janiszewska et al., 2017). Aragonitic blue coral fossils are also known from the Cretaceous (Eguchi, 1948; Colgan, 1984), further reinforcing the idea that the effect of different *m*Mg:*m*Ca ratios on the skeleton polymorph of corals differ between species and, likely, reflects the ability of some corals to deposit specific CaCO_3_ polymorphs under chemically unfavorable conditions.

*In vivo* experiments exposing animals to different *m*Mg:*m*Ca ratios provide support for the ability of corals to counteract the putative decisive influence of seawater chemistry on polymorph formation. In an early work, Ries (2006) observed an increase in the amount of calcite present in the skeleton of three scleractinian corals, with the amount of calcite found correlating inversely to the seawater *m*Mg:*m*Ca ratio. These observations were later corroborated by Higuchi et al (2014), who reported significant amounts of calcite in the skeleton of juvenile *Acropora tenuis*, albeit at lower *m*Mg:*m*Ca ratios compared to Ries (2006). More recently, Yuyama et al. (2019) reported changes in gene expression in the scleractinian coral *A. tenuis* grown in Mg-depleted seawater. These included the up-regulation of several putative skeletogenic genes, suggesting that lower *m*Mg:*m*Ca elicits a response in the corals’ calcification machinery. However, since Yuyama et al (2019) only manipulated the Mg-content of the water leaving the calcium levels constant, the molecular response of corals to *m*Mg:*m*Ca ratios comparable to those observed during geological periods corresponding to Calcite Seas, when both the concentration of calcium and magnesium changed, remain unknown.

Here, we exposed the branching stony coral *Montipora digitata* (Scleractinia) and the blue coral *Heliopora coerulea* (Octocorallia) to both calcite and aragonite-inducing seawater and used gene expression analysis (RNA-seq), electron backscatter diffractometry (EBSD), and energy dispersive spectroscopy (EDS) to investigate the molecular and mineralogical response of these corals to different *m*Mg:*m*Ca ratios resembling past oceanic conditions.

## Materials and Methods

Fragments of *H. coerulea* and *M. digitata* (ca. 1cm^3^) were mechanically obtained (and attached to the substrate) from colonies cultured for more than a decade in the aquarium facilities of the Chair for Geobiology & Palaeontology of the Department of Earth and Environmental Sciences at Ludwig-Maximilians-Universität München (Munich, Germany). Following fragmentation, corals were allowed to recover for about one month prior to the start of each experiment. Two days before the start of each experiment, coral samples were incubated for ca. 36 hours in a 10% alizarin red solution to mark the skeleton deposited before the experiment.

### Experimental Design

Two independent experiments were performed for each target species. Each experiment lasted ca. six weeks (22.07.-10.09.2018 and 01.11.-12.12.2018 for *M. digitata* and 04.01.-28.02.2019 and 01.03.-15.04.2019 for *H. coerulea)*. The experimental setup is illustrated in Figure 1. It consisted of three 8 L aquarium tanks: one (control tank) containing seawater with an *m*Mg:*m*Ca of 5.2 (10mM Ca^2+^ - 52 mM Mg^2+^) and two treatment tanks with ratios 2.5 (17mM Ca^2+^-47 mM Mg^2+^) and 1.5 (25mM Ca^2+^-37 mM Mg^2+^). In all three tanks, the sum of [Ca^2+^] and [Mg^2+^] was equal and set at 62 mmol. A single source of Mg^2+^ and Ca^2+^-free seawater was used for all three tanks and the *m*Mg:*m*Ca ratio was adjusted by adding different amounts CaCl_2_xH_2_O and MgCl_2_x6H_2_O to each tank. Water was replaced in each tank every two days. The composition of control and treatment artificial seawater used is provided in S.Mat1 (available online at https://github.com/PalMuc/CalciteSea).

**Figure 1.**
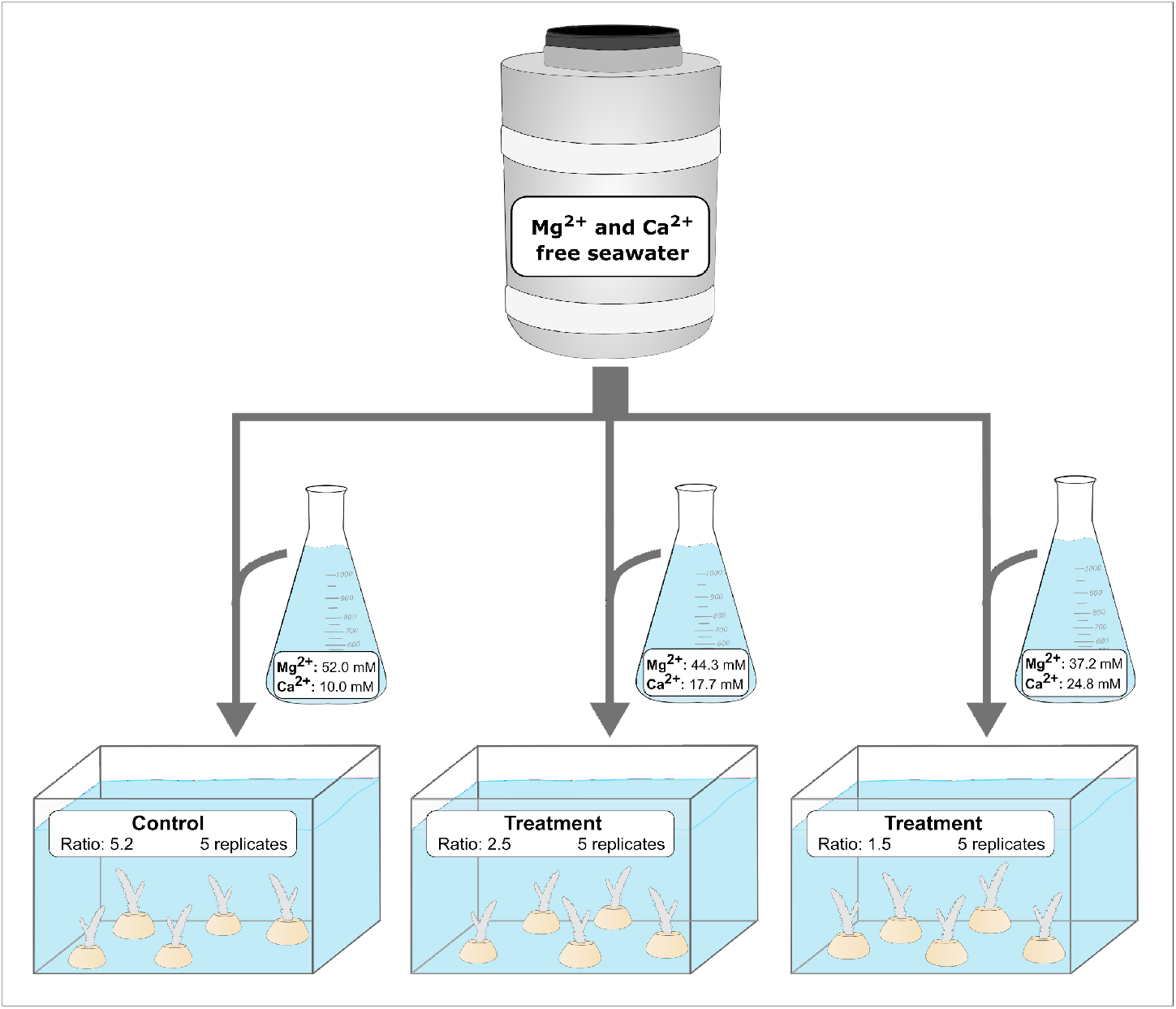
Schematic diagram of the experimental design. For each species the experiment was repeated twice: 22.07.-10.09.2018 and 01.11.-12.12.2018 for *M. digitata*, and 04.01.-28.02.2019 and 01.03.-15.04.2019 for *H. coerulea*.

To allow for acclimatization, corals in treatment tanks were initially exposed for two days to an *m*Mg:*m*Ca ratio of 5.2, before lowering the *m*Mg:*m*Ca ratio to 4.0 and 3.0 for two days, respectively. After the acclimation phase, the *m*Mg:*m*Ca ratio was adjusted to the final experimental ratios. At the end of each experiment, one coral fragment was bleached for 24 hrs in 5% NaOCl, washed several times in ultrapure water, air dried, and stored for mineralogical analysis. Three corals from each tank were flash-frozen in liquid nitrogen and stored at -80° C until RNA extraction.

### Mineralogical Analysis

To prepare coral samples for EBSD measurements, skeletons were first thoroughly washed with deionized water and then air dried. Samples were embedded in polypropylene moulds using ca. 30 ml of Epofix Resin (Struers) and 4 ml of Resin Hardener. Samples were placed in a vacuum desiccator to degas and polymerize for 24 hours. Embedded skeletons surfaces were ground using P320, P600, and P1200 silicon carbide paper (Buehler), and flushed with deionized water to remove resin and silicon residues. The first round of polishing was done with a 3 μm polycrystalline diamond suspension for ca. 10-15 minutes. The samples were then attached to cylinder weights with Thermoplastic Cement at 100-150°C. After reaching room temperature, samples were finally polished (for ca. three hours) with MicroFloc in a Vibromet using a 50 nm Alumina suspension and washed. Prior to EBSD analysis, the polished samples were coated with 4-8 nm of carbon and mounted on a sample holder with a 70° tilt with respect to the beam. For each sample, an ‘inner’ and ‘outer’ (growing edge) area were analyzed to obtain mineralogical information about the skeleton deposited before and after the start of the experiment. EBSD measurements were performed with a Hitachi SU5000 field emission SEM operated at 20kV. The microscope is equipped with an electron backscatter diffraction (EBSD) detector (Oxford Instruments). Measurements were conducted using the AZtech Suite (Oxford Instruments), while phase and orientation maps were produced with the CHANNEL 5 HKL software (Schmidt and Olesen, 1989; Randle and Engler, 2000). Electron backscatter diffraction measurements give crystal orientation and phase composition (Nowell and Wright, 2005). Crystal orientation is shown color-coded in EBSD. The used color code is shown with each EBSD map.

Grain parameters for all samples were determined with Tango 5.12. For each mapped skeleton portion, the relative abundance of grain area and major axis was computed. In an effort to account for differences possibly caused by sectioning plane and natural spatial heterogeneity of grain size, an additional variable “*minor axis”* was calculated by dividing each grain area by its major axis. For data plotting all three measures were log-transformed. Mineralogical datasets (both raw files and datasets with additional variables (minor axis)) - alongside the R script used to generate the grain statistics plots are available at https://github.com/PalMuc/CalciteSea. All SEM raw files used to generate phase images, band contrast images, and orientation maps have been deposited in the Open Data LMU repository (https://data.ub.uni-muenchen.de; DOI pending).

### Transcriptome Sequencing

Whole coral samples were homogenized in 1-2 ml of Trizol (Thermofisher) using a Polytron PT Homogenizer (Kinematica) and centrifuged at 15,000 g for 10 minutes to remove residual skeleton powder. RNA was extracted according to a modified TriZol protocol (Chomczynski and Mackey, 1995). Modifications included the substitution of 50% of the isopropanol with a high-salt solution (1.2 M sodium chloride and 0.8 M sodium citrate) in order to avoid the coprecipitation of polysaccharides and other potential contaminants. RNA purity and integrity were assessed on a NanoDrop 2100 spectrophotometer and a Bioanalyzer 2100 (Agilent), respectively. For sequencing, only samples with a RIN value > 8.0 were considered (N=17 *H. coeruela* and N=16 *M. digitata*). Strand-specific libraries were prepared with the SENSE mRNA-Seq Library Prep Kit V2 for Illumina (Lexogen) and paired-end sequenced (50 bp) on an Illumina HiSeq1500 at the Gene Center of Ludwig-Maximilians-Universität München. Sequences were quality controlled with FastQC (www.bioinformatics.babraham.ac.uk) and low-quality reads were removed with the Filter Illumina program from the Agalma-Biolite transcriptome package (Q value cutoff: 28) (Dunn et al., 2013). Reads generated as part of these experiments have been deposited at the European Nucleotide Archive (www.ebi.ac.uk/ena) under Bioproject Number PRJEB36989 (*M. digitata*) and PRJEB36990 (*H. coerulea*). For the assembly of *H. coerulea*, previously sequenced reads (PRJEB30452) were also used.

Transcriptomes were assembled with TransPi (Rivera-Vicéns et al., 2022). Contigs of length < 300 bp were discarded. Host and symbiont contigs were separated with psytrans (https://github.com/sylvainforet/psytrans) using the Dendronephthya gigantea (Jeon et al. 2019) and Acropora digitifera (Shinzato et al., 2011) sequences for assigning host contigs of H. coerulea and M. digitifera sequences, respectively. Transcriptome completeness was assessed with BUSCO 3.0.2 (metazoa database) (Simão et al., 2015). Summary statistics of the transcriptomes are provided in Table 1.

**Table.1.** Summary statistics for the *H. coerulea* and *M. digitata* transcriptomes assembled with Transpi. Parameters refer to host transcriptome CDS only after contigs assignment with psytrans. For BUSCO scores, percentages of complete (C), fragmented (F), and missing (M) are provided.

| Species | Contigs | N50 - Mean length | BUSCO [C-F-M] |
| --- | --- | --- | --- |
| *H. coerulea* | 110,535 | 819 - 716 | [ 97.0 - 1.4 - 1.6 ] |
| *M. digitata* | 71,676 | 819 - 705 | [ 90.4 - 4.3 - 5.3 ] |

### Gene Expression Analysis

For gene expression analysis, read files of each species were mapped against their metatranscriptome (host and symbiont contigs) using Salmon 0.11.2 (Patro et al., 2017) via our local Galaxy server (galaxy.palmuc.org). Read quantification files for each sample were combined into a count matrix with the abundance_estimates_to_matrix.pl script provided with Trinity 2.8.6 (Grabherr et al., 2011), and count data for host (coral) sequences was filtered based on psytrans results. Differential expression analysis was performed using DESeq2 v.1.24.0 (Love et al., 2014). To account for the observed batch effect between experimental replicates (Figures S5 and S6), the experimental replicate was included as a factor in the model for DESeq2 analysis using the design formula “*∼experiment + condition*”. For each comparison (i.e., *m*Mg:Ca 5.2 vs 2.5 and 5.2 vs 1.5), differentially expressed genes with adjusted p-value < 0.01 and log-fold changes - 2 > and > 2 were considered. The complete DESeq2 R-script workflow used, along with the count matrices, expression tables, sequences of differentially expressed genes, and the transcriptomic assemblies of *H. coerulea* and *M. digitata* are available as supplementary materials at https://github.com/PalMuc/CalciteSea.

## Results

### Mineralogical Analysis

EBSD measurements of *Heliopora coerulea* and *Montipora digitata* skeletons showed a possible species-specific response to variations in seawater *m*Mg:*m*Ca (Figures 2–4). Under control conditions (*m*Mg:*m*Ca = 5.2) and *m*Mg:*m*Ca of 2.5, calcite was not detected in both *H. coerulea* or *M. digitata* (Figure 3–4). This pattern was consistent between the two experimental replicates. Contrary, the presence of calcite was detected in one *M. digitata* coral sample maintained under calcite-inducing conditions (*m*Mg:*m*Ca of 1.5) (Figure 2), whereas only aragonite was observed in the *H. coerulea* skeleton areas surveyed under these conditions.

**Figure 2a:**
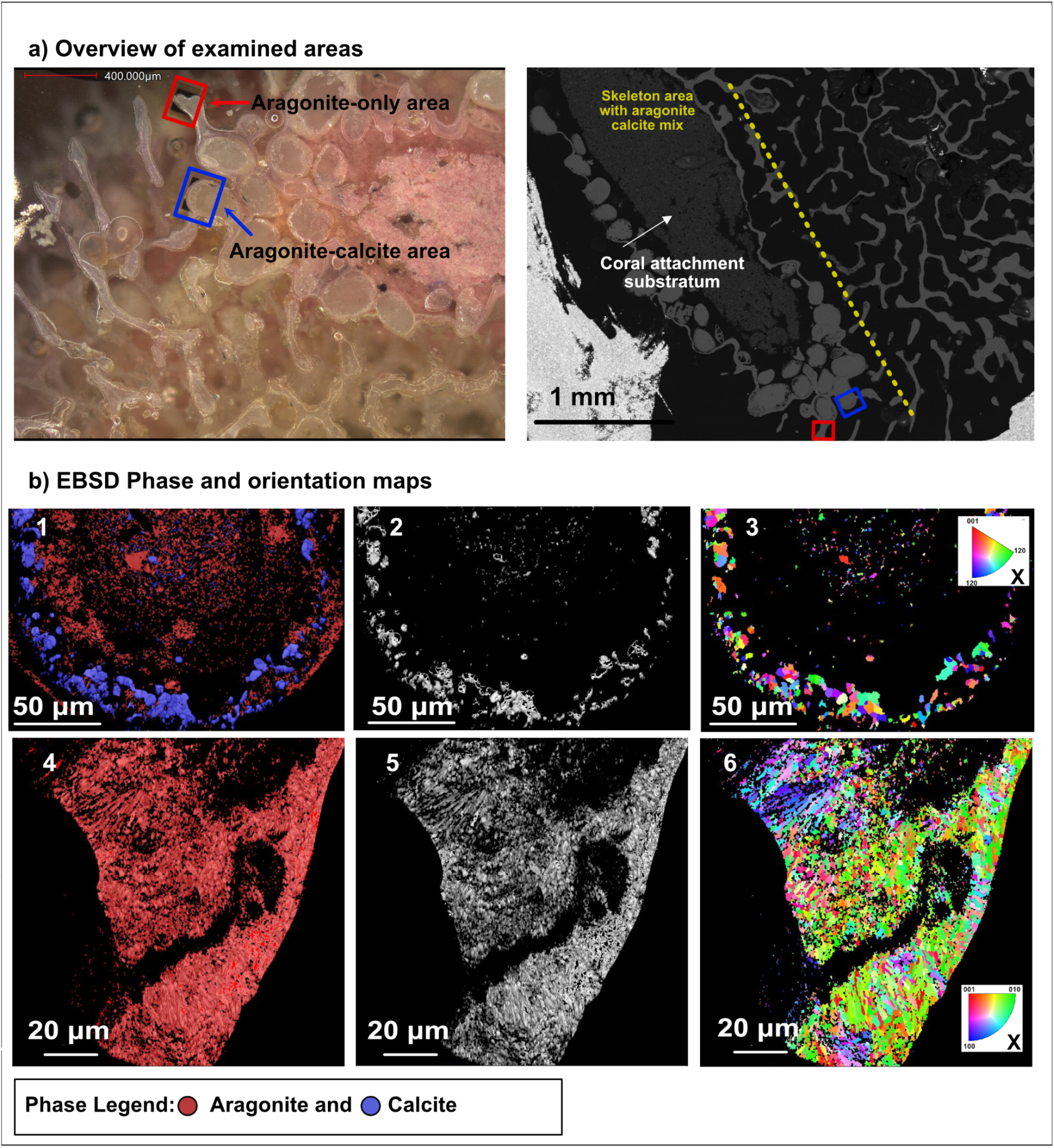
Light microscopy (left) and SEM (right) image of *Montipora digitata* skeleton with calcite. Red and blue squares indicate two areas deposited during the experiment and analyzed with EBSD. Data for the area deposited prior to the experiment is available in Figure S1 **b)** Phase map, band contrast, and orientation map of the area where calcite was observed (1-3) (blue rectangle) and the area composed only of aragonite (4-6) (red rectangle).

**Figure 3.**
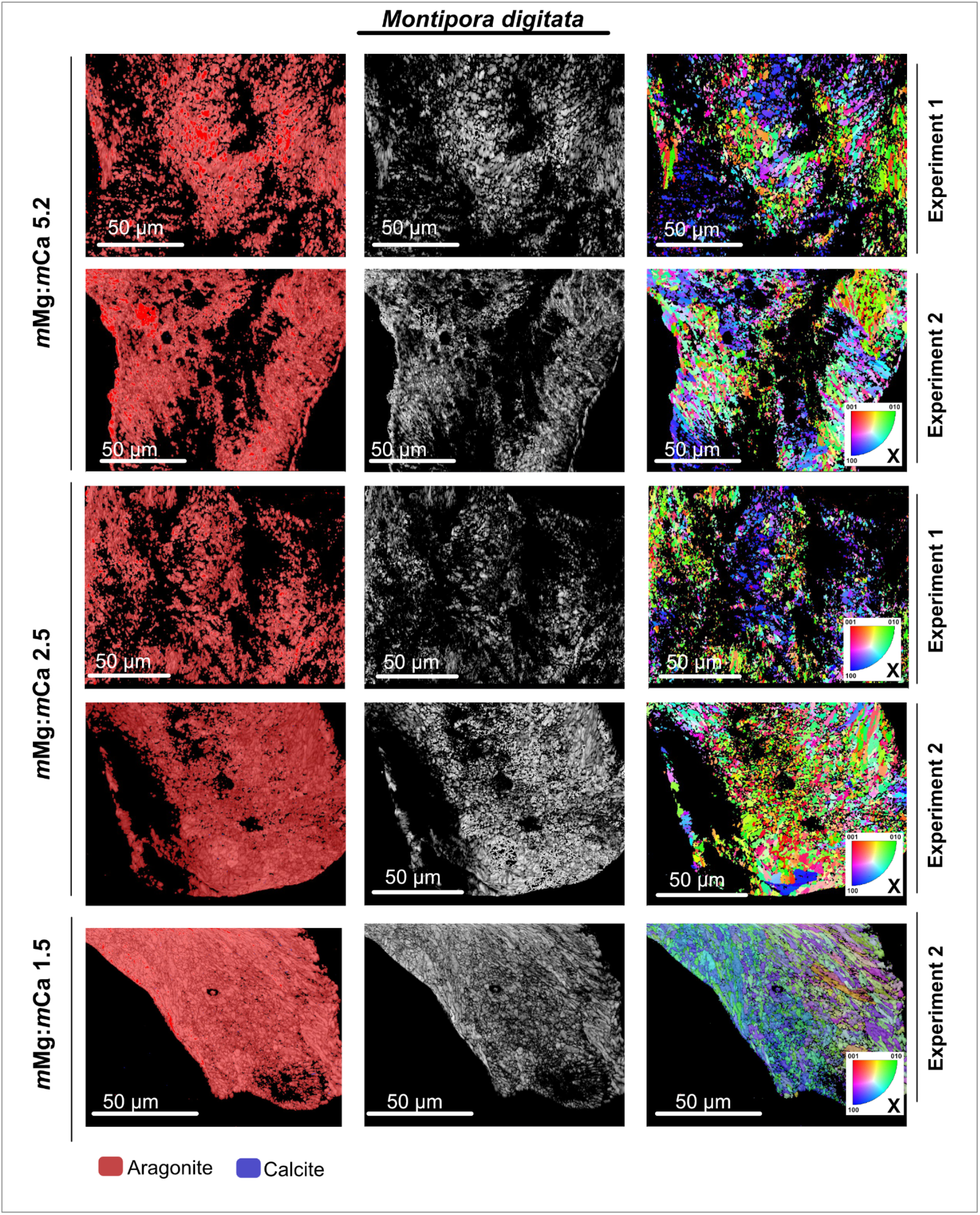
Calcium carbonate phase map, band contrast images, and EBSD orientation map of *Montipora digitata* skeletons grown under each experimental condition (*m*Mg:*m*Ca 5.2, 2.5, and 1.5) and during experimental replicate. Information for the *M. digitata* sample in which calcite was observed is available in Figure 2. In the phase map figures, red indicates aragonite and blue indicates calcite.

Microscopy images (Figure 3a, 3b) show that the calcite did not form homogeneously within the newly deposited skeleton, but it was rather found to be restricted within mineralized circular structures. These appear in direct contact with the coral skeleton. Phase analysis (Figure 4 b1) revealed a mixed composition within these areas with the co-presence of both aragonite and calcite. Neither these circular structures nor calcite could be detected in the portion of the skeleton grown prior to the experiment (Figure S1). Analysis of grain area and diameters showed differences between species, experimental conditions, and replicates (Figures S3 and S4). In *M. digitata*, no difference in terms of grain area or both major and minor grain axes in corals exposed to an *m*Mg:*m*Ca of 2.5, while in one coral grown at an *m*Mg:*m*Ca of 1.5 a decrease in grain size - compared to control conditions - could be observed (Figure S3 and S4). On the contrary, in one experiment replicate both corals grown under treatment conditions (2.5 and 1.5) exhibited larger grain sizes (Figures S3 and S4). For both species, differences between control and treatment ratios were larger when grain areas were compared, while a higher overlap could be observed when grain minor axes were compared.

**Figure 4.**
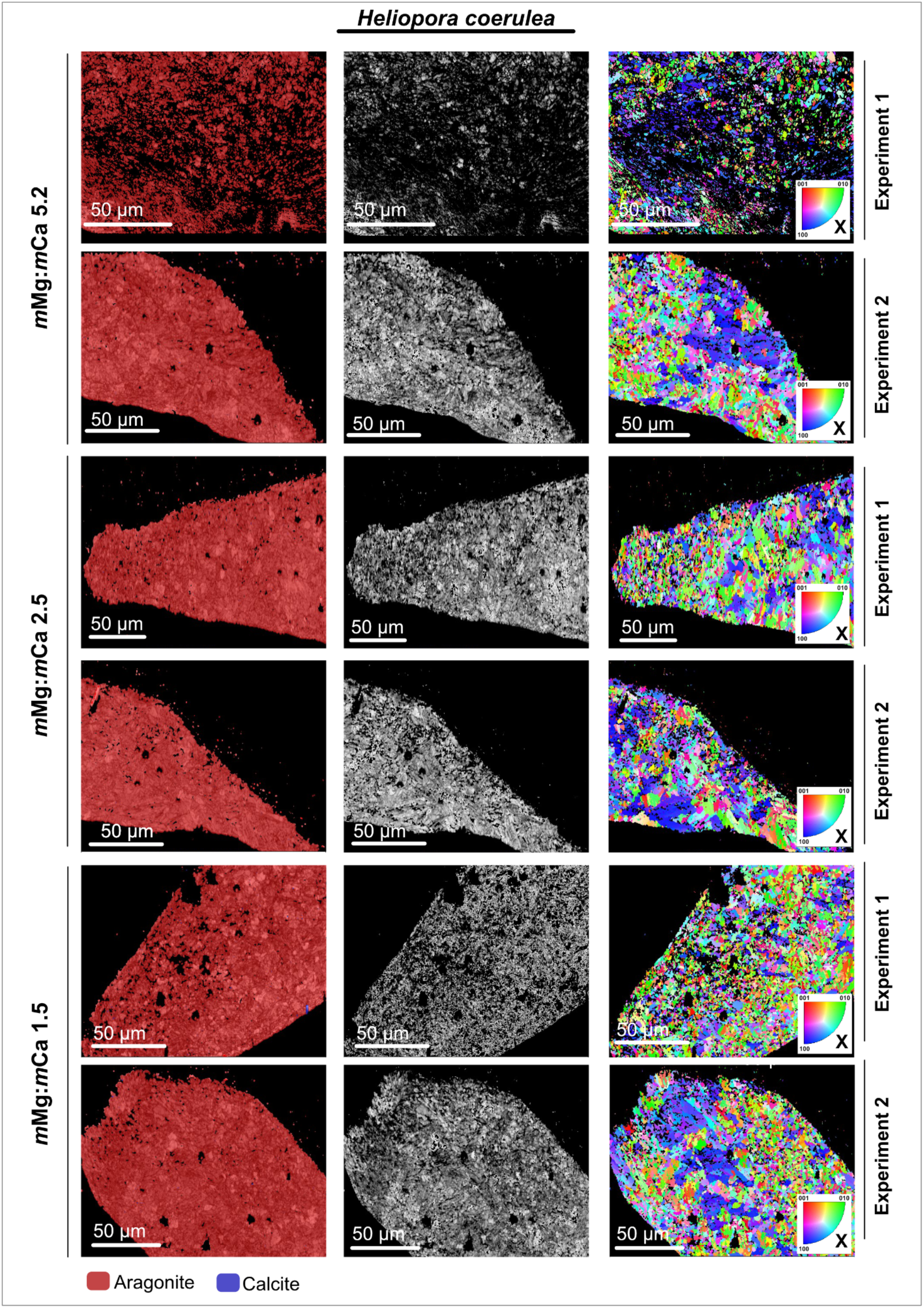
Calcium carbonate phase map, band contrast images, and EBSD orientation map of *Heliopora coerulea* skeletons grown under each experimental condition (*m*Mg:*m*Ca 5.2, 2.5 and 1.5) and experimental replicate. In the phase map figures, red indicates aragonite and blue indicates calcite.

### Gene Expression Analysis

Exposure to *m*Mg:*m*Ca ratios of 2.5 and 1.5 triggered changes in gene expression (p-value < 0.001; log fold change > |2|) in both species. The octocoral *H. coeruela* showed overall higher numbers of differentially expressed genes (DEGs) compared to *M. digitata* under both treatments. However, principal component analysis (PCA), based on log-transformed expression values relative to control, revealed a high dispersion of control and treatment samples for both species, and strong differences between the two experimental replicates for both species (Figures S5 and S6). In *M. digitata*, expression changes of known putative calcification-related genes were restricted to *m*Mg:*m*Ca of 1.5 (Figure 5). These included the over-expression of one galaxin-like 2 (AD50285.1) homolog, one matrix metalloproteinase-like protein, and one uncharacterized skeletal matrix protein (B3EX00.1) previously identified in the skeleton of scleractinian corals (Ramos-Silva et al., 2013; Conci et al., 2020). In *H. coerulea*, the only putative calcification DEG found coded for a cartilage-intermediate layer like-protein and was consistently down-regulated in blue corals exposed to *m*Mg:*m*Ca of 2.5.

**Fig 5:**
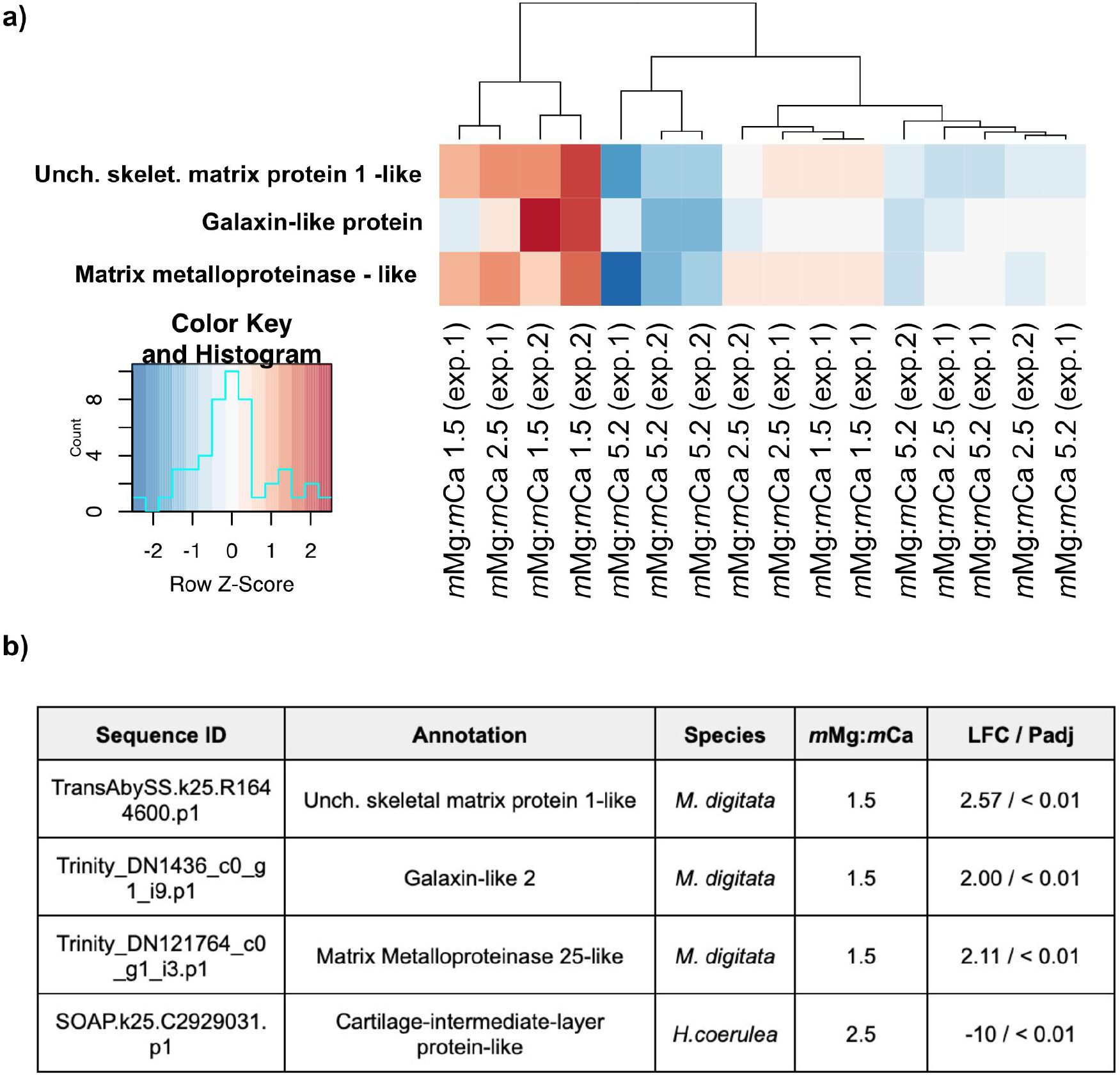
**a)** Gene expression heatmap of potential calcification-related genes in *M. digitata* under different *m*Mg:*m*Ca ratios. **b)** Table providing accession IDs, annotation, and log fold-change (LFC) and adjusted p-value (Padj). The table also includes the only potential calcification-related gene we observe changing across experiments in *H. coerulea*.

Analysis of single experimental replicates showed up-regulation for additional potential calcification-related genes. In *M. digitata*, these included, coadhesin-like and protocadherin-like reported in the *A. millepor*a and *S. pistillata* skeletons (Ramos-Silva et al., 2013; Drake et al., 2013). Moreover, in one experimental replicate, galaxin was also up-regulated at *m*Mg:*m*Ca of 2.5. Finally, an uncharacterized skeletal organic matrix protein 5-like protein was found down-regulated at *m*Mg:*m*Ca of 1.5.

Although changes in the expression of calcium or magnesium transporters were not detected in both experimental replicates, different voltage-dependent calcium channel alpha subunits were down-regulated in *H. coerulea* under *m*Mg:*m*Ca of 2.5 in Experiment 1.

## Discussion

The formation of coral skeletons is intimately linked to both cellular/molecular processes, and the surrounding environmental conditions. Among the latter, the magnesium/calcium molar ratio of seawater (*m*Mg:*m*Ca) appears to have influenced the polymorph of calcium carbonate (CaCO_3_) of major marine calcifiers, and the polymorph initially adopted by marine organisms during the last 500 myr (Zhuravlev and Wood, 2008). However, organisms also appear to be less affected by subsequent changes in seawater chemistry promoting the precipitation of a different polymorph (Porter, 2010). Manipulative experiments conducted in artificially recreated “Calcite Seas” have, however, highlighted the inability of modern scleractinian corals to fully counteract the effects of *m*Mg:*m*Ca changes, leading to significant amounts of calcite being deposited within their skeletons (Ries et al., 2006; Higuchi et al., 2014). The present study combined mineralogical (electron backscattered diffraction, EBSD) and molecular (gene expression) analyses to provide a first insight into the effect of different Mg:Ca molar ratios on aragonitic octocoral skeletons, namely those of *Heliopora coerulea*, and compared the results with the scleractinian coral *Montipora digitata*. To examine and compare their responses at the polymorph and gene expression level, both species were exposed to three different *m*Mg:*m*Ca ratios: the modern value of 5.2 (control group), 2.5 and 1.5.

As previously shown for scleractinian species (Ries et al., 2006; Higuchi et al., 2014), our experiments highlighted the ability of *M. digitata* and *H. coerulea* to deposit aragonite when exposed to calcite-inducing *m*Mg:*m*Ca ratios down to 1.5. Albeit not being detected in both samples, the presence of calcite in the *M. digitata* samples also supports the observations from the two aforementioned studies. Although we could not detect calcite in corals gown at *m*Mg:*m*Ca ratios higher than 2, as previously described in *M. digitata* <u>(Ries et al., 2006)</u>

In this regard, differences in the experimental setup, analytical methods, and both coral target species, as well as growth stages have to be considered when comparing results across different studies. Although the characterization of the CaCO_3_ polymorph with EBSD allowed us to obtain information about crystal size and orientation, it was here restricted to delimited areas of a limited number of coral colonies. This might 1) represent one of the reasons for the variability observed between experimental replicates for several of the parameters investigated, and 2) have possibly caused to have missed and/or underestimated calcite content in the skeletons. Future research should thus include additional methodologies (e.g., X-ray diffraction or Raman spectroscopy) to more comprehensively assess the presence of polymorph changes and allow the investigation of higher sample numbers.

The origin and formation mechanisms of the calcite observed at low *m*Mg:*m*Ca ratios in *M*.*digitata* remain elusive. The distribution analysis of the detected calcite grains points to the calcite-inducing effects of the *m*Mg:*m*Ca ratio not being limited to polymorph substitutions (same structure but different composition). On the contrary, calcite was only observed in areas that also exhibit distinct macro-morphological differences. Compared to the rest of the skeleton – which exhibits branching-like structures – calcite in *M. digitata* appears as spherical bodies surrounded by (and potentially in direct contact with) the aragonitic skeleton.

In this regard, although the observed morphological variation could be related to the effects of *m*Mg:*m*Ca on the biological calcification machinery, we also consider the possibility that the observed structures originate in enclosed areas of *M. digitata*’s skeleton when corals are exposed to calcite-inducing conditions. In these chambers, the composition of calcification fluids might experience strong shifts and promote the precipitation of calcite alongside aragonite. Finally, it cannot be excluded that the presence of calcite could also be possibly related to other mechanisms, such as the random presence of foreign mineral material, that has been previously observed in scleractinian skeletons (Goffredo et al., 2012).

Variability was observed between experiments aiming to investigate the molecular mechanisms behind the effects of *m*Mg:*m*Ca on coral calcification. Consistent across experiments, expression data from *H. coerulea* highlighted expression changes for only one putative calcification-related gene: one cartilage-intermediate layer protein homolog underexpressed under *m*Mg:*m*Ca of 2.5. No expression changes for components of the known skeleton proteome of this species (Conci et al., 2020) were detected. On the other hand, in *M. digitata*, a set of calcification-related genes, including a galaxin-related protein, was found upregulated in both experimental replicates, when the *m*Mg:*m*Ca was lowered to 1.5. The response of SkOM genes to calcite-inducing *m*Mg:*m*Ca described here might nonetheless have been underestimated, as the high variability observed between and within experiments might have prevented the detection of changes in the expression of additional genes. Overexpression of multiple SkOM proteins (including galaxin-related proteins) under calcite-inducing *m*Mg:*m*Ca has nonetheless been previously reported in *A. tenuis* (grown under *m*Mg:*m*Ca of 0.5), and was linked to active initiation and maintenance of aragonite deposition under adverse environmental conditions (Yuyama and Higuchi, 2019). Generally, the overexpression of calcification-related genes in corals exposed to calcite-inducing environments may indicate an increased energetic expenditure by the corals and explain the decline in calcification rates observed in different coral species at lower *m*Mg:*m*Ca ratios (Higuchi et al., 2014; Ries et al., 2006). The reduction in grain sizes observed here in *M. digitata* may also represent a consequence of a higher energetic cost of calcification under low mMg:mCa ratios. However, as for the gene expression analyses, the level of variation observed across experiments and possible effects of biological variability and sample preparation on the measured grain parameters prompt the need for additional research.

In addition to SkOM components, variation in *m*Mg:*m*Ca ratios could also potentially affect ion transport and/or chemistry of the calcification fluids. Although inconsistent between experiments, observed changes in the expression of calcium transporters in *H. coerulea* could be related to a response of the ion transport machinery used by corals for calcification. An involvement of calcium transporters has been excluded for *A. tenuis*, as no change in expression was detected (Yuyama and Higuchi, 2019). However, in that study the seawater *m*Mg:*m*Ca was manipulated by decreasing [Mg^2+^], while keeping [Ca^2+^] at normal levels (ca. 10mM). Thus, information on the putative response of calcium and other ion transporters in corals concurrently exposed to low [Mg^2+^] and increased [Ca^2+^] is currently lacking. In corals, calcium is supposedly transported to the calcification sites either paracellularly (Gagnon et al., 2012) or via active transcellular transport (Tambutté et al., 1996; Hohn and Merico, 2015). Different transporters have been associated with transcellular calcium transport including calcium channels, Ca^2+^-ATPases, and proton exchangers (Zoccola et al., 1999, 2004; Barott et al., 2015; Barron et al., 2018). The regulation of calcium transport allows corals to maintain higher calcium concentrations in calcifying fluids ([Ca^2+^]_cf_) with respect to seawater (Sevilgen et al., 2019). Up-regulation of [Ca^2+^]_cf_ can also be enhanced in some coral species to counteract decreases in the saturation state of the calcifying medium caused by decreases in seawater pH (DeCarlo et al., 2018). Different studies have in fact examined the effects of ocean acidification on carbonate and calcium chemistry at calcification sites (Allison et al., 2018; Mollica et al., 2018). However, to the best of our knowledge, research examining the response of ion transport to the calcification site when the environmental calcium concentration is higher than [Ca^2+^]_cf_ is currently lacking. Further characterizations of the calcification-related calcium transport machinery in octocorals and scleractinians could provide additional insights into the localization and role of single calcium channels, and possibly corroborate the hypothesis that corals might control aragonite precipitation, when exposed to high environmental [Ca^2+^], through the regulation of calcium transport.

Here, we examined possible changes in skeleton polymorph and in the expression of calcification-related genes in the aragonitic octocoral *H. coerulea* and the scleractinian *M. digitata* exposed to calcite-inducing *m*Mg:*m*Ca. This work provides the first comparative study of octocoral and scleractinian responses to changes in *m*Mg:*m*Ca. The two species were both found to continue producing aragonite, while the presence of calcite could only be detected in the skeleton of *M. digitata*. Despite variability both within and between experiments, the observed changes in gene expression support previously hypothesized molecular responses of SkOM genes (Yuyama and Higuchi, 2019) when the *m*Mg:*m*Ca is lowered.

## Acknowledgments

The study was supported by the German Research Foundation (DFG) grant “MINORCA” to S.V. (Project Number Va1146-2/1) and G.W. (Project Number Wo896/18-1). G.W. acknowledges support by LMU Munich’s Institutional Strategy LMUexcellent within the framework of the German Excellence Initiative. We thank Peter Naumann for technical assistance and maintenance of the aquaria facilities and coral culturing, as well as Simone Schätzle and Gabriele Büttner for assistance in the laboratory and Moritz Zenkert for EBSD samples preparation.

## Notes

https://github.com/PalMuc/CalciteSea

### Competing Interest Statement

The authors have declared no competing interest.

### Summary of Updates

revised after comments

