## Supplementary Figure for "Molecular and mineral responses of corals grown under artificial Calcite Sea conditions"

### Supplementary Figures

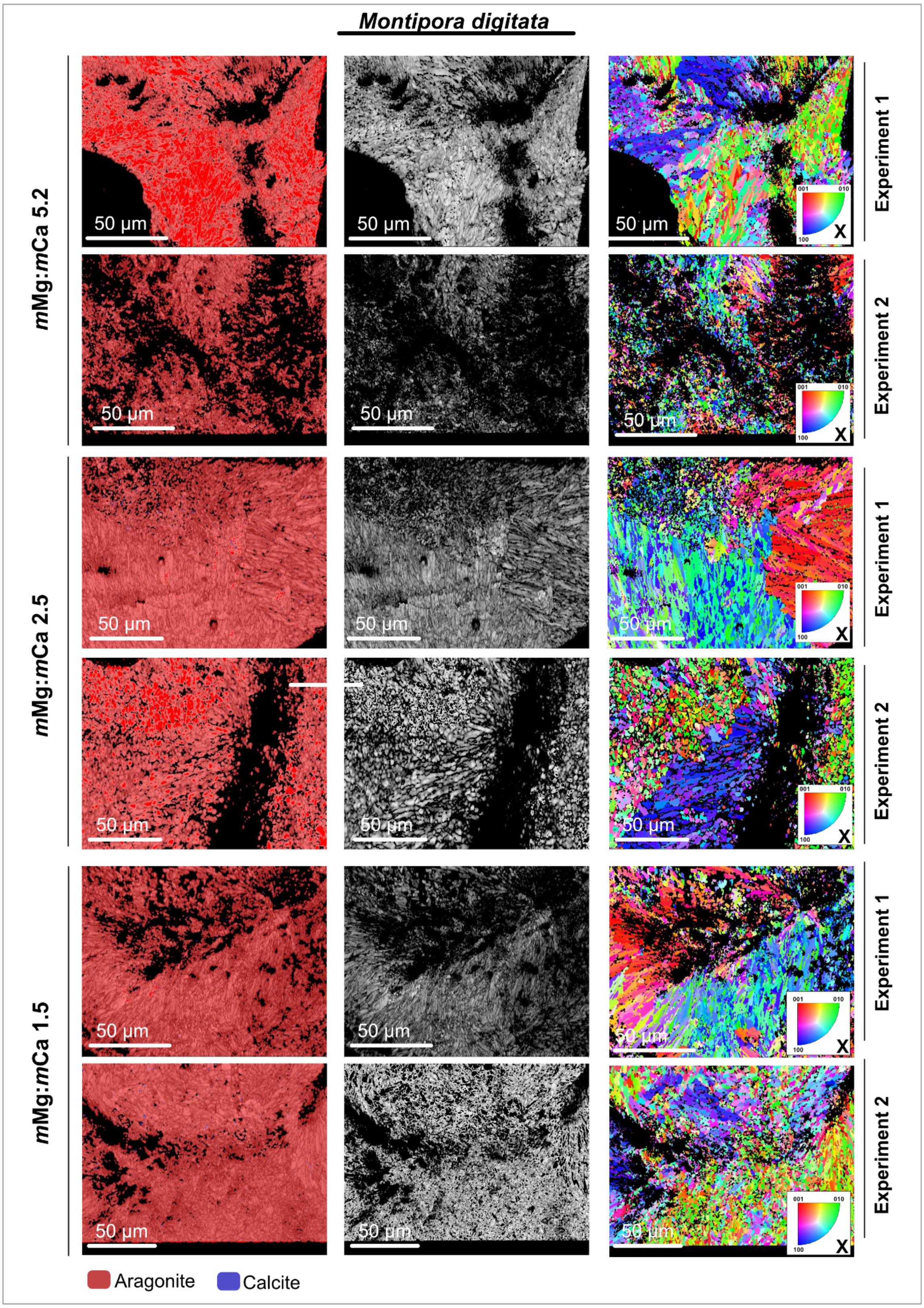


**Figure S1** Calcium carbonate phase map, band contrast images, and EBSD orientation map of *Montipora digitata* skeletons grown before each experiment. In phase map figures red indicates aragonite and blue indicates calcite.

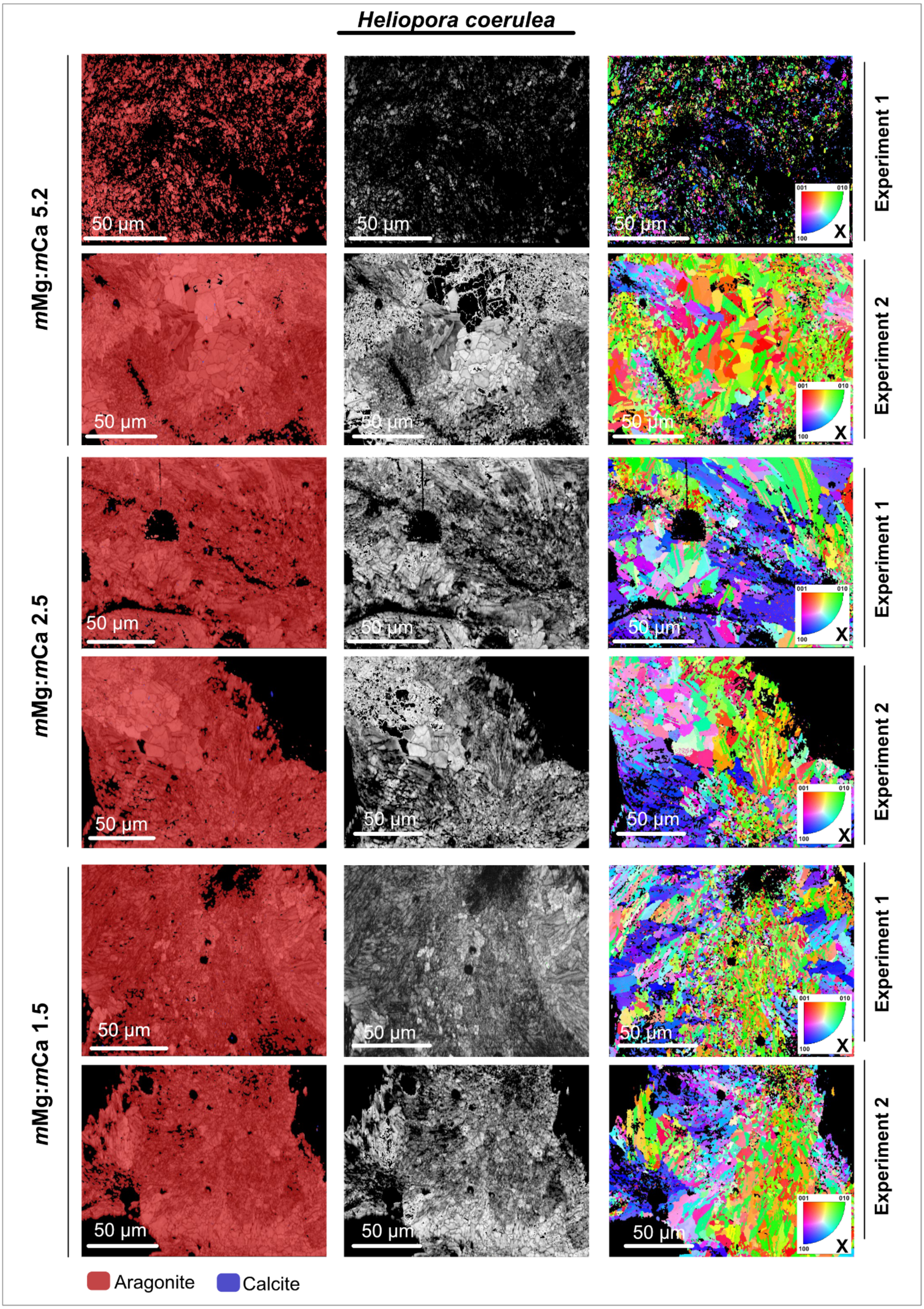


**Figure S2** Calcium carbonate phase map, band contrast images, and EBSD orientation map of *Heliopora coerulea* skeletons grown before each experiment. In phase map figures red indicates aragonite and blue indicates calcite.

### Grains statistics *Montipora digitata*

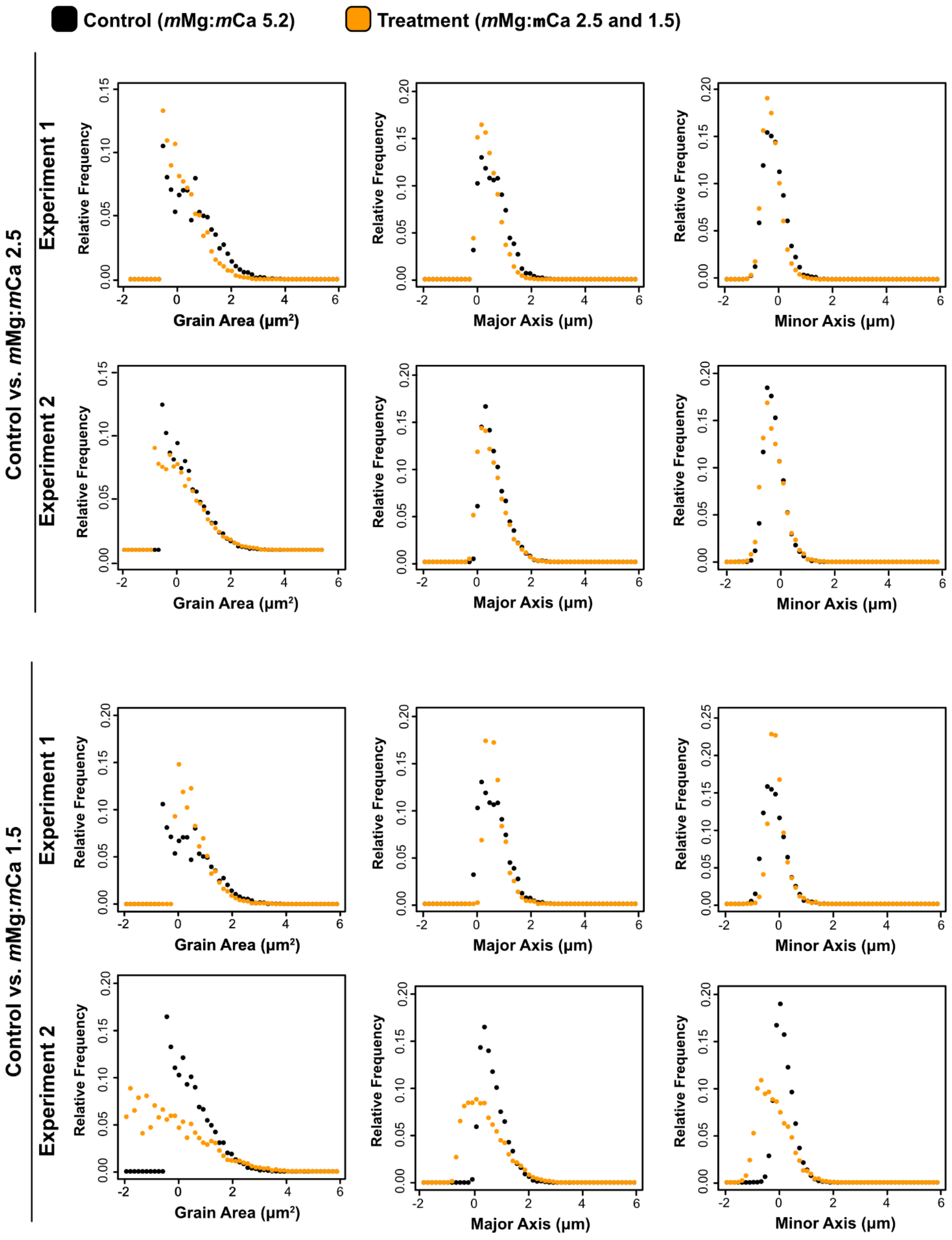


**Figure S3:** Frequency distribution of grain area, major and minor axis (log-transformed) in *Montipora digitata* corals grown under different *m*Mg:*m*Ca during both experimental replicates. Datasets and script to generate plots available as supplementary at https://github.com/PalMuc/CalciteSea.

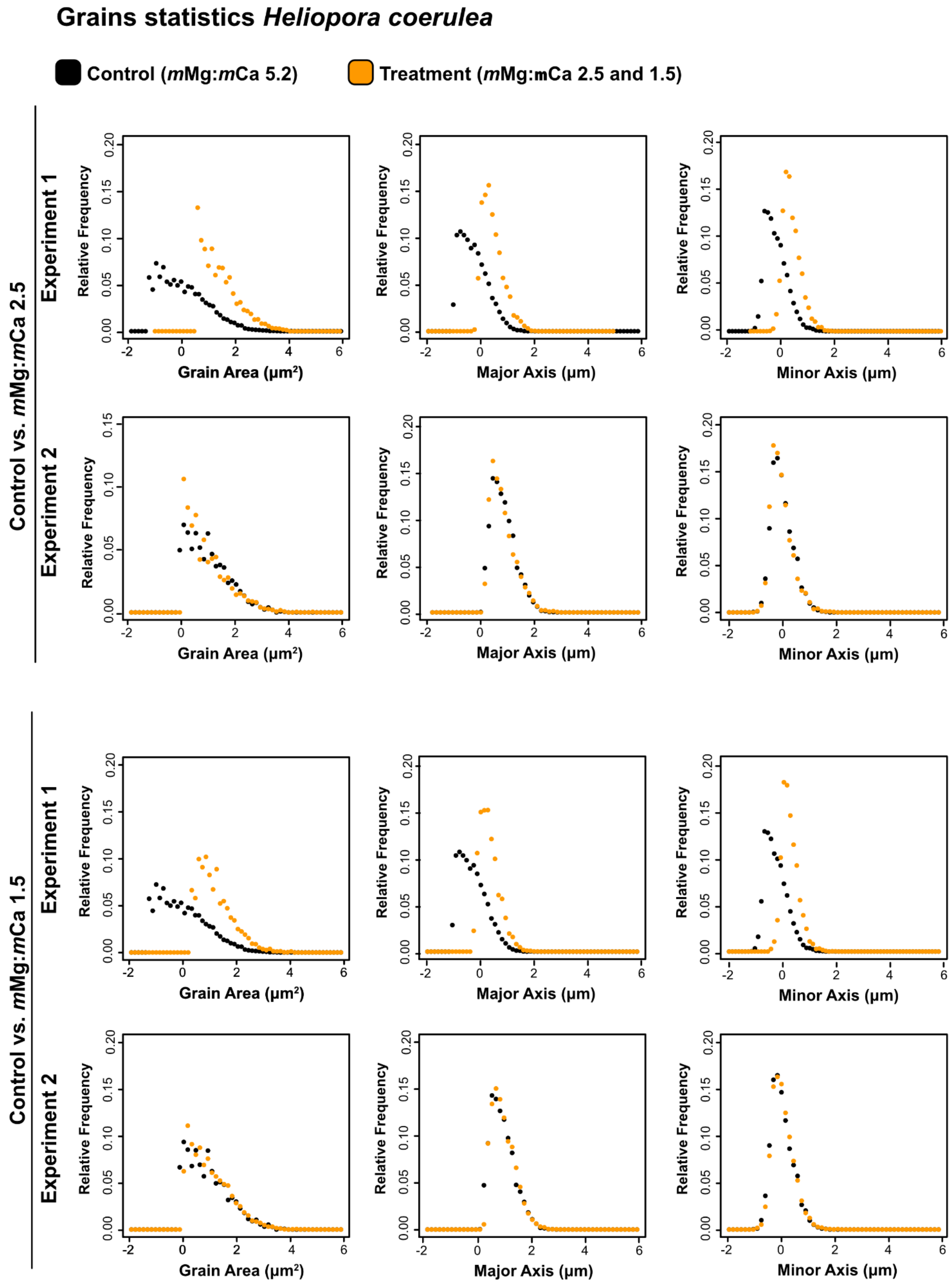


**Figure S4:** Frequency distribution of grain area, major and minor axis (log-transformed) in *Heliopora coerulea* corals grown under different *m*Mg:*m*Ca during both experimental replicates. Datasets and script to generate plots available as supplementary at https://github.com/PalMuc/CalciteSea.

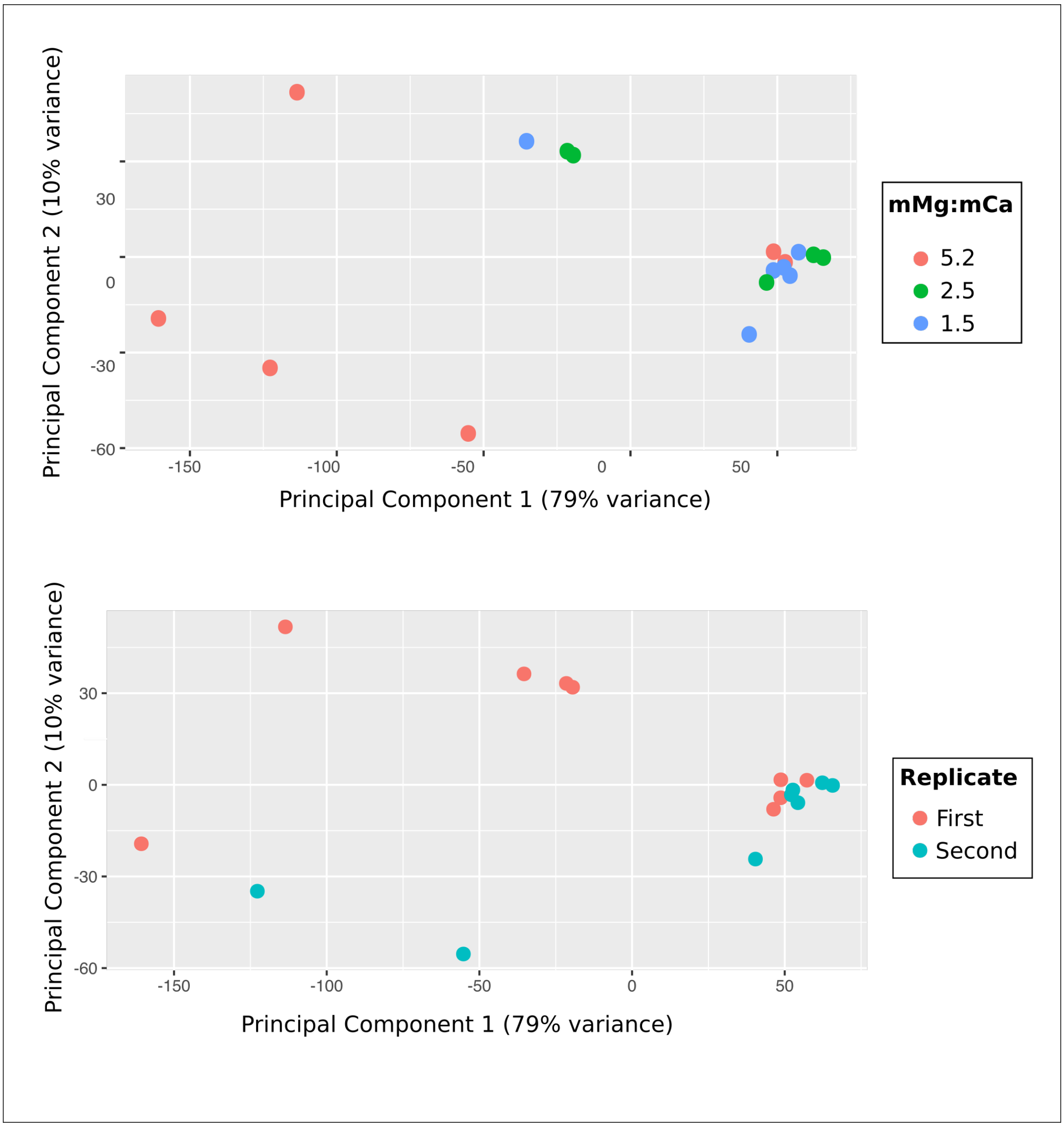


**Figure S5:** Principal component analysis for expression of genes assigned by psytrans to *M. digitata*. Dots are coloured based on experimental conditions (*m*Mg:*m*Ca) (top) and experimental replication (bottom).

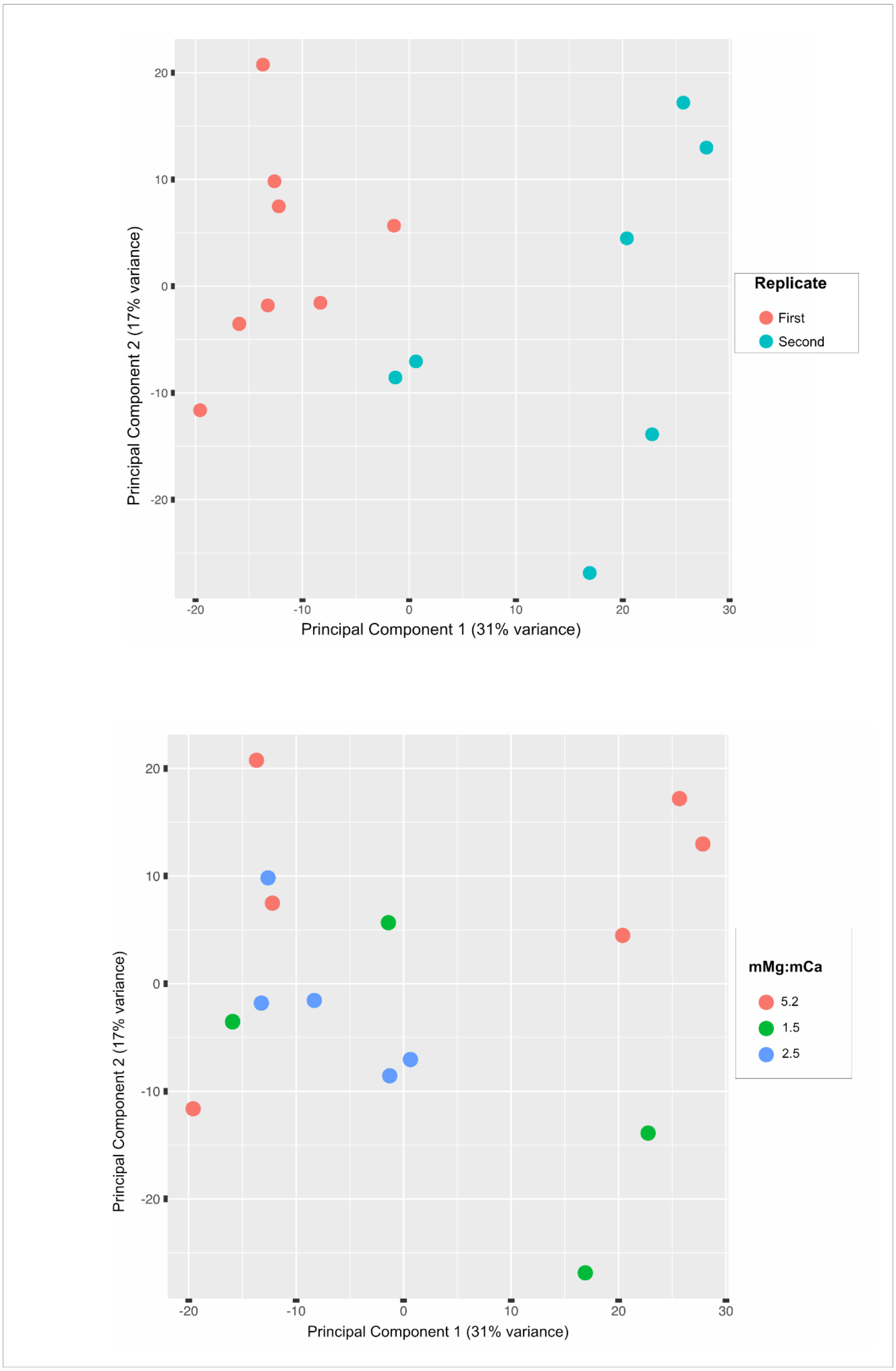


**Figure S6:** Principal component analysis for expression of genes assigned by psytrans to *M. digitata*. Dots are coloured based on experimental conditions (*m*Mg:*m*Ca) (top) and experimental replication (bottom).
